# Unsupervised selection of optimal single-molecule time series idealization criterion

**DOI:** 10.1101/2021.02.07.430124

**Authors:** Argha Bandyopadhyay, Marcel P. Goldschen-Ohm

## Abstract

Single-molecule (SM) approaches have provided valuable mechanistic information on many biophysical systems. As technological advances lead to ever-larger datasets, tools for rapid analysis and identification of molecules exhibiting the behavior of interest are increasingly important. In many cases the underlying mechanism is unknown, making unsupervised techniques desirable. The Divisive Segmentation and Clustering (DISC) algorithm is one such unsupervised method that idealizes noisy SM time series much faster than computationally intensive approaches without sacrificing accuracy. However, DISC relies on a user selected objective criterion (OC) to guide its estimation of the ideal time series. Here, we explore how different OCs affect DISC’s performance for data typical of SM fluorescence imaging experiments. We find that OCs differing in their penalty for model complexity each optimize DISC’s performance for time series with different properties such as signal-to-noise and number of sample points. Using a machine learning approach, we generate a decision boundary that allows unsupervised selection of OC based on the input time series to maximize performance for different types of data. This is particularly relevant for SM fluorescence datasets which often have signal-to-noise near the derived decision boundary and include time series of nonuniform length due to stochastic bleaching. Our approach allows unsupervised per-molecule optimization of DISC, which will substantially assist rapid analysis of high-throughput single-molecule datasets with noisy samples and nonuniform time windows.

## Introduction

Recent advances in detector and imaging technologies have enabled increasingly high-throughput SM data collection (Chen et al., 2014; Soltani et al., 2014; Yan et al., 2018). For example, faster sCMOS cameras with larger chips coupled with photobleach-resistant fluorophores allow simultaneous recording of hundreds of molecules per field of view for extended durations (Altman et al., 2012; Halabi et al., 2019; Juette et al., 2016). Tools for rapid unsupervised analysis of such large datasets are increasingly critical to avoid becoming the bottleneck for experiment progress (Miller et al., 2018; Xiaobo Zhou & Wong, 2006).

Hidden Markov models (HMMs) are a widely successful approach for SM time series analysis (Blanco & Walter, 2010; Bronson et al., 2009; McKinney et al., 2006). However, HMMs are computationally expensive, making analysis of large datasets an extremely time-consuming process that can take days to weeks on a typical high-end desktop computer (Blanco et al., 2013; White et al., 2020). Furthermore, HMMs require postulation of a molecular mechanism (i.e. specification of a model’s discrete set of states and the allowed transitions between them), which is often unknown a priori (Blanco & Walter, 2010). Although methods exist for selecting a model from a set of HMMs (Greenfeld et al., 2012), this poses a heavy computational burden proportional to the size of the test model set. Unsupervised approaches for HMM model selection (e.g. infinite HMMs (Hines et al., 2015; Sgouralis & Pressé, 2017) and deep learning neural networks (Celik et al., 2020; Li et al., 2020; Xu et al., 2019) automate the process of model identification, but remain computationally expensive or require extensive training datasets prior to their use.

To handle the scale of high-throughput datasets, efficient unsupervised approaches that do not require postulation of a specific mechanism or pre-training on similar known datasets allow rapid screening of the data which can be an important guide for choice of experiment conditions. Even if a more computationally expensive approach such as an HMM is ultimately desired, unsupervised approaches that can rapidly screen large datasets to identify the subset of molecules displaying the behavior of interest are of enormous value. In most experimental conditions, only a subset of the observed molecules is of interest for further analysis. Performing costly computations on huge numbers of molecules that are ultimately not included in the final analysis is incredibly inefficient. In many cases, selection of a subset of molecules of interest is performed manually, which is both time consuming and biased. Thus, fast preliminary analysis with an unsupervised method is desirable for the majority of SM datasets irrespective of whether or not it is the final analysis.

The Divisive Segmentation and Clustering (DISC) algorithm is an unsupervised top-down approach for rapid idealization of noisy piecewise continuous time series typical of SM imaging experiments (White et al., 2020). For a given noisy time series, DISC estimates the underlying ideal noiseless time series consisting of discrete jumps between a finite number of distinct intensity levels. Both the jumps and the number of distinct intensity levels is determined in an unsupervised fashion and does not require the user to postulate a molecular mechanism prior to analysis. Compared to HMMs or change point analyses, DISC is orders of magnitude faster while maintaining state-of-the-art accuracy, precision and recall. Lately, many deep learning techniques reliant on neural networks have been developed for unsupervised SM analysis (Celik et al., 2020; Li et al., 2020). Unlike these approaches, DISC does not require extensive training datasets to guide its idealization which simplifies its application to multiple different experimental regimes. Rather than relying on training data, DISC utilizes a user specified objective criterion (OC) that weighs goodness of fit against the complexity of the ideal sequence to guide idealization (Kadane & Lazar, 2004). The OC represents a metric for unsupervised approaches to identify the simplest model that yet describes the noisy experimental data reasonably well, while avoiding complex models that overfit the data. In addition to DISC, OCs have been widely applied in numerous SM analysis approaches including HMM model selection and change point idealization (Bronson et al., 2009; Greenfeld et al., 2012; McKinney et al., 2006; Shuang et al., 2014).

Here, we show that different OCs optimize DISC’s accuracy, precision and recall for SM time series that differ in their signal-to-noise ratio or number of sample points. This is crucial for SM fluorescence imaging experiments where nonuniformity in the optics or illumination and stochastic bleaching of fluorophores results in variable SNRs and observation window durations across molecules even within a single field of view. To maximize the performance of DISC on such a dataset, we use a machine learning technique to automate the per-molecule selection of the optimal OC. Critically, this automation makes DISC robust to both the scale and heterogeneity of datasets typical of increasingly common high-throughput SM imaging experiments (Holden et al., 2010; Oostveldt et al., 1998).

## Results

### Different OCs optimize DISC’s performance under different experimental conditions

The DISC algorithm is an unsupervised top-down approach for rapid idealization of noisy piecewise continuous time series typical of SM imaging experiments (White et al., 2020). Although subsequently unsupervised, the algorithm requires a user-selected OC to guide its idealization. DISC has three main steps: 1) Divisive segmentation: Starting with the intensity values of all data points in the time series assigned to a single cluster (the mean intensity of the time series), each cluster is recursively split into two child clusters until the selected OC is met. A major advantage of this approach is that typically only a few levels of segmentation are needed. Thus, the following agglomeration step is much faster than typical agglomerative approaches that start with each data point in its own individual cluster. 2) Hierarchical agglomerative clustering: During segmentation, clusters in separate branches that should belong to the same intensity level may be assigned unique intensity states due to random fluctuations in the data. Agglomerative clustering starting with the segmented clusters from the previous step remerges segments with similar intensity distributions based on the selected OC. 3) Viterbi: The previous steps accurately identify intensity levels in the time series, but do not provide a good description of the kinetics of transitions amongst those levels. To estimate event kinetics, the Viterbi algorithm is applied to determine the most likely sequence of transitions amongst the identified intensity levels. We have previously shown that DISC provides orders of magnitude faster computational speed while maintaining comparable accuracy, precision and recall to state-of-the-art approaches (White et al., 2020). However, the impact of OC choice on the accuracy of DISC’s idealization for different kinds of data has yet to be rigorously evaluated.

To explore the impact of OC choice on DISC’s performance, we simulated noisy SM time series with known state sequences and evaluated the ability of DISC to identify the correct noiseless sequences using different OCs. We simulated data for several different kinds of mechanisms to evaluate a variety of typical SM data. Furthermore, we varied simulation parameters such as sample length, signal to noise, and state transition rates to determine the impact of OC choice on data under different experimental conditions. SM time series were simulated for several mechanisms including two-state dynamics at one, two or four independent sites similar to binding observations from colocalization SM fluorescence experiments, and three-state linear or cyclic models with distinct state emissions typical of smFRET experiments **(Figure 1-figure supplement 1)**. Time series were simulated with uniform time steps and Gaussian noise for variable numbers of samples per trace (10^2^ to 10^4^), signal-to-noise ratios (SNRs; 1 to 8), and state transition rates (0.01 to 1 transitions per time step, on average) (see Methods). Here, we define SNR as the ratio of the intensity level separation to the standard deviation of the noise fluctuations. In many cases real experimental SM fluorescence data additionally contains heterogeneity in state emissions (Goldschen-Ohm et al., 2016; White et al., 2020). The source of this heterogeneity is uncertain but is likely caused by shifts of the molecule in the exponentially decaying excitation field or dye photodynamics (Dempsey et al., 2009; Levene et al., 2003). Changes in observed dye brightness due to dye conformation, polarization orientation, partial quenching via electron transfer and protein-induced fluorescence enhancement are commonly observed (Stennett et al., 2015). To better reflect these real observations, we additionally included heterogeneous state intensities in some of our simulations (see Methods).

Five OCs were tested: Bayesian information criterion assuming either a Gaussian mixture model (BIC_GMM_) or residual sum of squared errors (BIC_RSS_), Akaike information criterion (AIC), Hannan-Quinn information criterion (HQC), and minimum description length (MDL) (Eqs. 2-10) (Akaike, 1974; Hannan & Quinn, 1979; Priestley, 2004; Schwarz, 1978; Shuang et al., 2014). In each case, DISC’s idealization performance on the noisy simulated data was evaluated using standard criteria of accuracy, precision, and recall (Eqs. 11-13). An overall measure of this performance is summarized in the F1 score (0-1: worst to perfect) which combines both precision and recall in a single metric (Eq. 14) (Van Rijsbergen, 1979). For each OC, DISC’s performance was primarily dependent on the SNR (i.e. intensity level separation vs noise) and number of samples in the time series **(Figure 1-figure supplements 2-8)**. State transition rates had relatively less of an effect on performance except where rates were extreme (e.g. average rate equal to the sample rate). For data with heterogeneous state intensities there was no single OC that was optimal for all simulated conditions. Thus, individual time series conditions dictate the optimal choice of OC. However, for nearly all conditions the optimal OC was either BIC_GMM_ or BIC_RSS_ **(Figure 1-figure supplements 2-8)**.

In general, BIC_RSS_ outperforms BIC_GMM_ for short traces with less than about 1000 samples and low SNRs of ∼3 or less **(Figure 1 *top left*)**, whereas BIC_GMM_ outperforms BIC_RSS_ for longer traces with more samples and/or higher SNR **(Figure 1 *top right*)**. This is largely because BIC_GMM_ tends to underfit shorter series with low SNR whereas BIC_RSS_ overfits longer series with high SNR. Each OC balances a goodness of fit term and a penalty term that attempts to prevent overfitting for overly complex models/sequences (Eq. 1). The observed behavior can be understood by examination of the penalty terms for BIC_GMM_ and BIC_RSS_ (Eqs. 2-3). Because the number of level transitions or change points is a stochastic function of the length of the time series, BIC_RSS_ will generally have a lower penalty than BIC_GMM_ for shorter traces with few transitions. For short traces with a low SNR, BIC_GMM_ is likely to underfit the number of distinct intensity levels due to the small amount of data and relatively large variation around each level, whereas the relatively smaller penalty for adding a level with BIC_RSS_ means that such underfitting is less likely **(Figure 1 *top left*)**. However, the lower penalty for additional levels also means that BIC_RSS_ tends to split levels with heterogeneous event intensities into multiple sublevels due to a marginal increase in transitions-per-sublevel compared to a significant reduction in the sum of squared residuals. The amount of heterogeneity in each level will increase with the number of transitions into each level, which increases with the length of the series. Also, as SNR increases such heterogeneity becomes more distinct. Thus, for longer traces with a high SNR, heterogeneous event intensities are likely to be overfit by BIC_RSS_ in comparison to BIC_GMM_ **(Figure 1 *top right*)**. In the absence of heterogeneous state intensities, BIC_RSS_ either matches or outperforms BIC_GMM_ for most experimental conditions. However, BIC_GMM_ continues to outperform BIC_RSS_ for longer traces with a moderate SNR (between 3 and 7) at rapid transition rates (e.g. average transition rate approaching sampling rate) **(Figure 1-figure supplements 7-8)**. This, again, can be understood by examination of the penalty terms for each OC (Eqs. 2 and 3). At exceptionally fast rates, long traces will have large numbers of change points which result in a larger penalty term for BIC_RSS_ than for BIC_GMM_. Due to this larger penalty, BIC_RSS_ is more prone to underfit these traces than BIC_GMM_.

**Figure 1.**
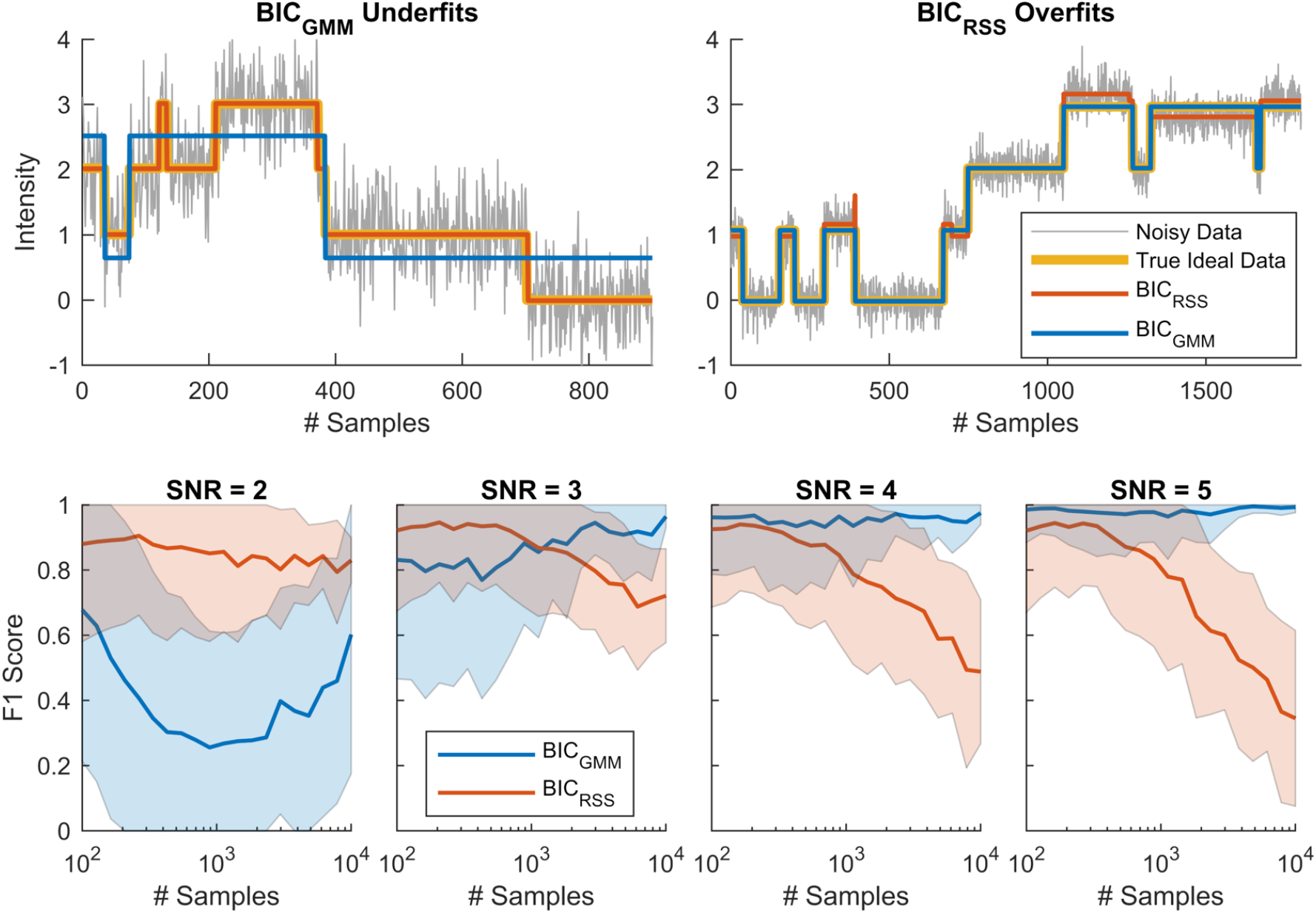
Optimal choice of BIC_GMM_ versus BIC_RSS_ depends on experimental conditions. ***top*)** Examples of simulated SM time series for the 4-site model in **Figure 1-figure supplement 1** with and without added noise. Traces are overlaid with DISC’s idealization of the noisy data using either BIC_GMM_ or BIC_RSS_. BIC_GMM_ tends to result in underfitting of short time series with less than about 1000 samples and lower signal-to-noise ratios (SNR) (*top left*), whereas BIC_RSS_ tends to result in overfitting of longer time series with higher SNR (*top right*). ***bottom*)** Summary of performance for DISC’s idealization of simulated noisy SM time series across a range of series lengths and several SNRs. Mean (line) and standard deviation (shaded region) for F1 scores (0-1: worst to perfect; Eq. 14) for 10-1000 simulated time series at each unique set of conditions (# samples and SNR) idealized with DISC using either BIC_GMM_ or BIC_RSS_ (see Methods). See **Figure 1-figure supplements 2-8** for additional conditions and models.

Therefore, optimal idealization with DISC requires selection of either BIC_GMM_ or BIC_RSS_ based on the number of samples and SNR in the time series for each individual molecule. For SM fluorescence measurements where fluorophore bleaching and spatial or temporal heterogeneity in excitation power can lead to stochastic variability in both the number of samples and SNR across molecules, this choice should optimally be made on a per-molecule basis.

### Optimization of a variable penalty hyperparameter does not outperform either BIC_GMM_ or BIC_RSS_

Given the dependence of DISC’s performance on the penalty term of the chosen OC, we explored whether optimizing a variable penalty term would further enhance performance. We introduced a hyperparameter λ that scales the penalty term (Eq. 15). Using either BIC_GMM_ or BIC_RSS_ as the framework for the OC, we scaled λ from 0.001 to 10 and chose the value of λ which maximized performance for a given set of experimental conditions (i.e. number of samples and SNR) **(Figure 1-figure supplements 9-10)**. This markedly improved performance for each OC in their respective problematic regimes. However, in nearly all conditions this approach did not outperform the better of either BIC_GMM_ or BIC_RSS_. Thus, a simple selection between BIC_GMM_ or BIC_RSS_ is sufficient for optimal performance on a per-molecule basis.

### A decision boundary for selecting the optimal OC on a per-molecule basis

To automate the unsupervised optimal choice of either BIC_GMM_ or BIC_RSS_ on a per-molecule basis, we used a machine learning tool, the support vector machine (SVM), to identify a two-dimensional linear decision boundary based on number of samples and SNR in each time series **(Figure 2)**. For each unique pair of conditions (number of samples and SNR), **Figure 2** illustrates the fraction of 4-site model simulations where performance was optimal for either BIC_GMM_ or BIC_RSS_. The boundary between optimal OCs is well described by a line, and thus more complex nonlinear boundaries were unnecessary. Constraining the boundary to a linear SVM also prevents overfitting small fluctuations in the training dataset.

**Figure 2.**
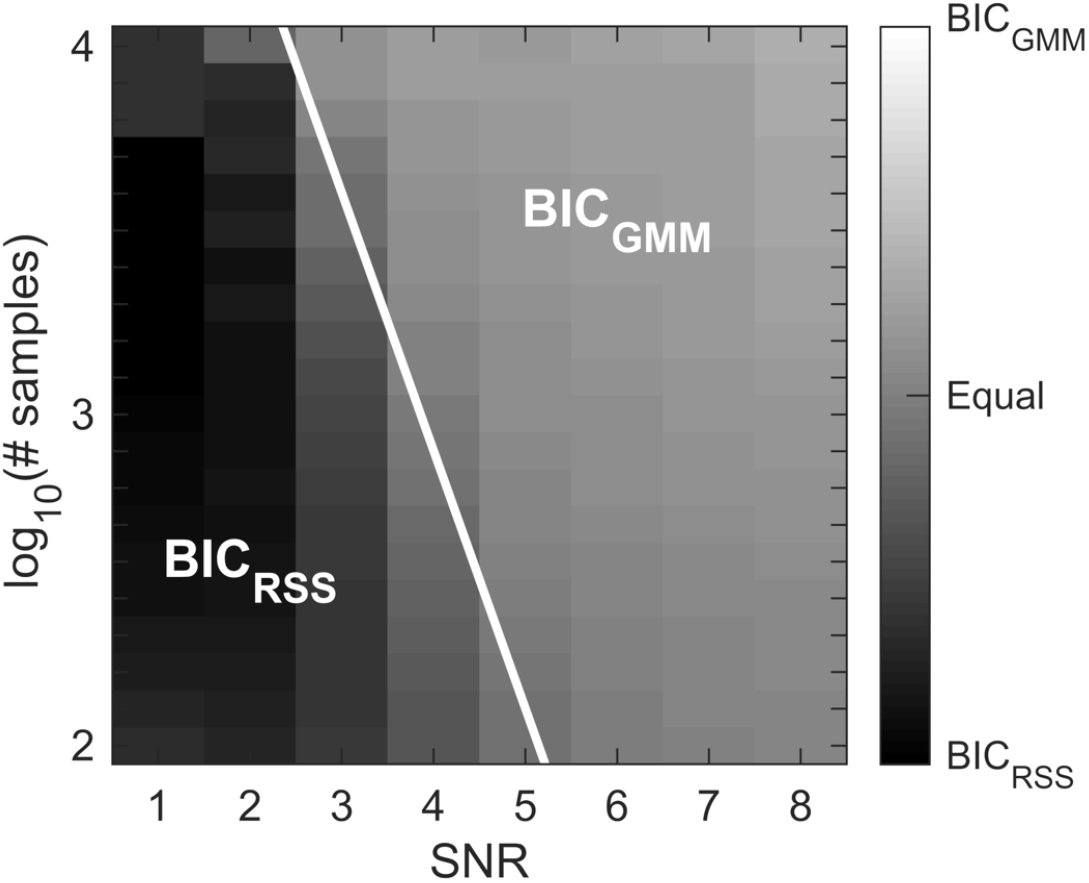
A linear decision boundary for the optimal choice of either BIC_GMM_ or BIC_RSS_. Heatmap of the degree of preference for BIC_GMM_ versus BIC_RSS_ (see Eq. 16) for 4-site model simulations with state intensity heterogeneity. Preference was determined from 1000 simulated time series for each unique pair of conditions (SNR and number of samples in the time series). The white line denotes the linear decision boundary determined by the SVM classifier.

A similar linear decision boundary was determined for each model in **Figure 1-figure supplement 1**. The boundaries for each model were largely similar but differed slightly in their slope and intercept, with the primary difference being a shift of about ±1-2 in SNR **(Figure 2-figure supplements 1-2)**. In all cases, the region within about +2-3 SNR to the right of the identified decision boundary (i.e. higher SNR) is a region where both BIC_GMM_ and BIC_RSS_ perform nearly equally well. This “buffer” region contains an infinite family of roughly equally appropriate decision boundaries. Notably, the variation in decision boundary across models was smaller than the buffer region, such that a single boundary appropriate for all models could be readily identified. Here, we selected the boundary for the 4-site model as appropriate for all tested models. This boundary is given by *log*_10_(#*samples*) =−0.74 *SNR* + 5.8. We cannot rule out that some mechanisms may give rise to time series with very different decision boundaries. However, the set of models and range of simulated conditions covers representative data for typical SM fluorescence experiments. Thus, the identified boundary is likely to be relevant for many SM imaging datasets or similar data.

### Estimation of SNR for SM time series

To use this decision boundary, one must estimate the separation in intensity levels and the noise of each time series in an experimental dataset. Therefore, we developed an unsupervised approach to estimate the average SNR of an SM time series **(Figure 3)**. First, we apply DISC using BIC_RSS_ to generate an initial idealization that may overfit, but likely does not underfit the data. We then estimate the noise from the residuals after subtracting this idealization from the raw data. To estimate the average separation between intensity levels in the time series, we first cluster the step heights of all intensity changes in the idealized series into two groups using binary *k*-means. The reason for this clustering is that in the case where the initial idealization overfits the data, the lower amplitude cluster will reflect overfitting of noise fluctuations, and thus can be discarded so as not to distort our estimation of the true level separations. The mean of the higher amplitude cluster is then taken as an estimate of the average level separation in the series. For series with nonuniform intensity level separations, this will be a weighted average based on the frequency of transitions between levels. Finally, the SNR is estimated as the ratio of the average level separation to the standard deviation of the residuals.

**Figure 3.**
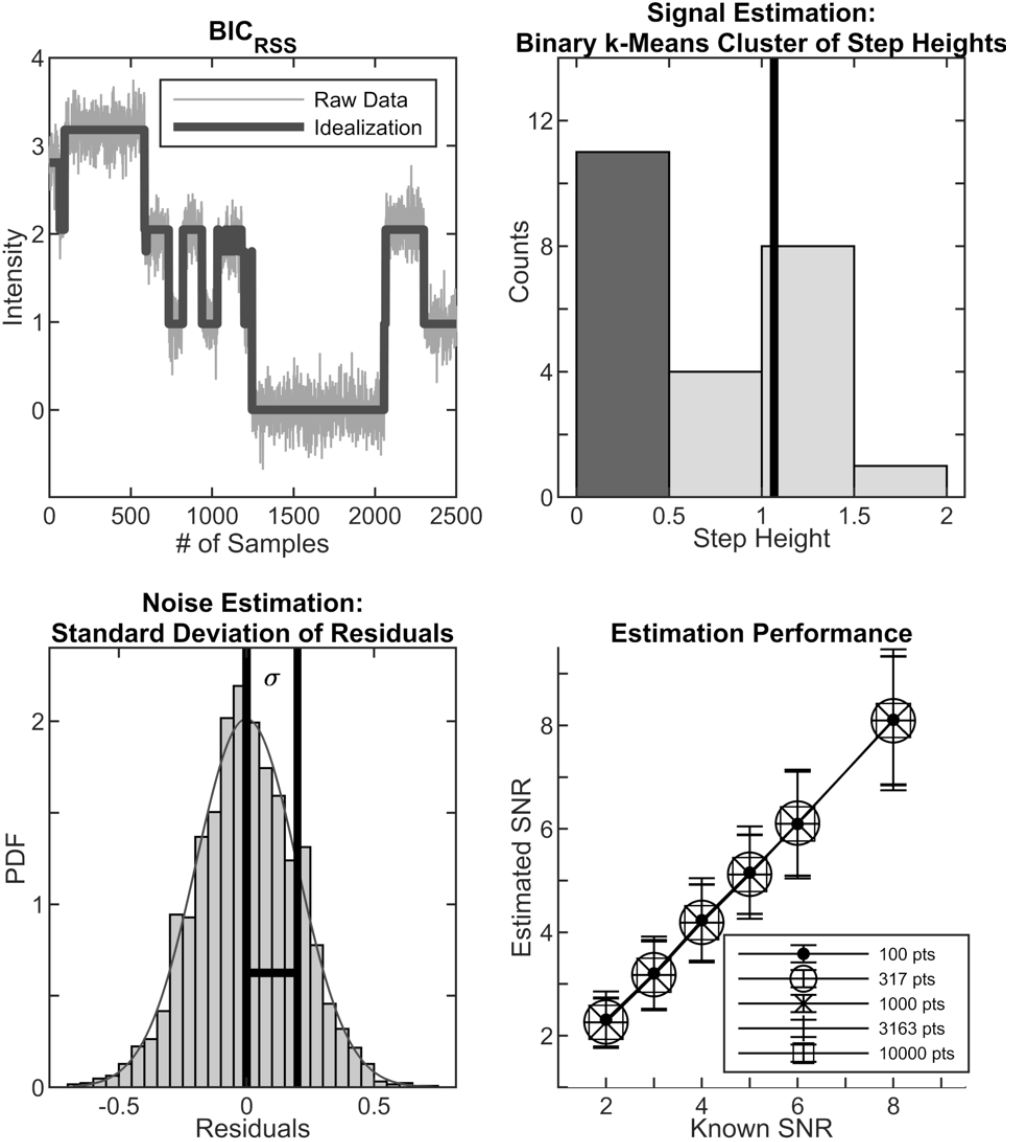
Estimation of the average intensity level separation and noise in a time series. *top-left*) Idealization of a noisy time series using DISC with BIC_RSS_. *top-right*) Histogram of step heights from the idealized series to the left shaded based on their classification into two clusters via binary *k*-means. The mean of the larger amplitude cluster (vertical line) is taken as the estimate for the average intensity level separation in the time series. *bottom-left*) Histogram of the residuals after subtracting the idealization from the noisy time series above. The standard deviation of the distribution of residuals is taken as the estimate for the average noise in the observations. Residuals were normally distributed, as shown by the overlaid Gaussian fit. *bottom-right*) Estimated versus known SNR for simulated time series (10-1000 series totaling 100,000 samples) under various conditions (simulated SNR and number of sample points per trace) for the four independent site model with average transition rate one tenth of the sample rate and added state intensity heterogeneity. SNR is computed as the ratio of the average intensity level separation to noise within each time series.

We tested the SNR estimation approach described above on each of the models in **Figure 1-figure supplement 1**. For all models, the estimation was highly accurate except for cases where transition rates were extreme (e.g. equivalent to the sample rate). However, in the case of extremely fast transition rates both BIC_GMM_ and BIC_RSS_ perform roughly equally well **(Figure 1-figure supplements 2-8)**, such that an erroneous estimation of SNR is unlikely to result in an inappropriate choice of OC.

### A completely unsupervised workflow for optimal per-molecule performance with DISC

With the linear decision boundary for optimal choice of either BIC_GMM_ or BIC_RSS_ and the SNR estimation approach described above, we can now establish a completely unsupervised workflow for optimizing the DISC algorithm on a per-molecule basis **(Figure 4)**. The workflow for an individual time series is as follows: 1) Apply DISC with BIC_RSS_ to idealize the time series. 2) Estimate the SNR of the series. 3) Based on the number of samples and the estimated SNR, select the optimal OC based on the linear decision boundary between BIC_GMM_ and BIC_RSS_. 4) If BIC_GMM_ is optimal, apply DISC with BIC_GMM_ to idealize the time series. Otherwise use the initial idealization. This workflow allows unsupervised per-molecule idealization of noisy SM data with stochastic variation in series duration, SNR and event intensities typical of many SM fluorescence imaging datasets. Software implementing this workflow is available at https://github.com/marcel-goldschen-ohm/DISC.

**Figure 4.**
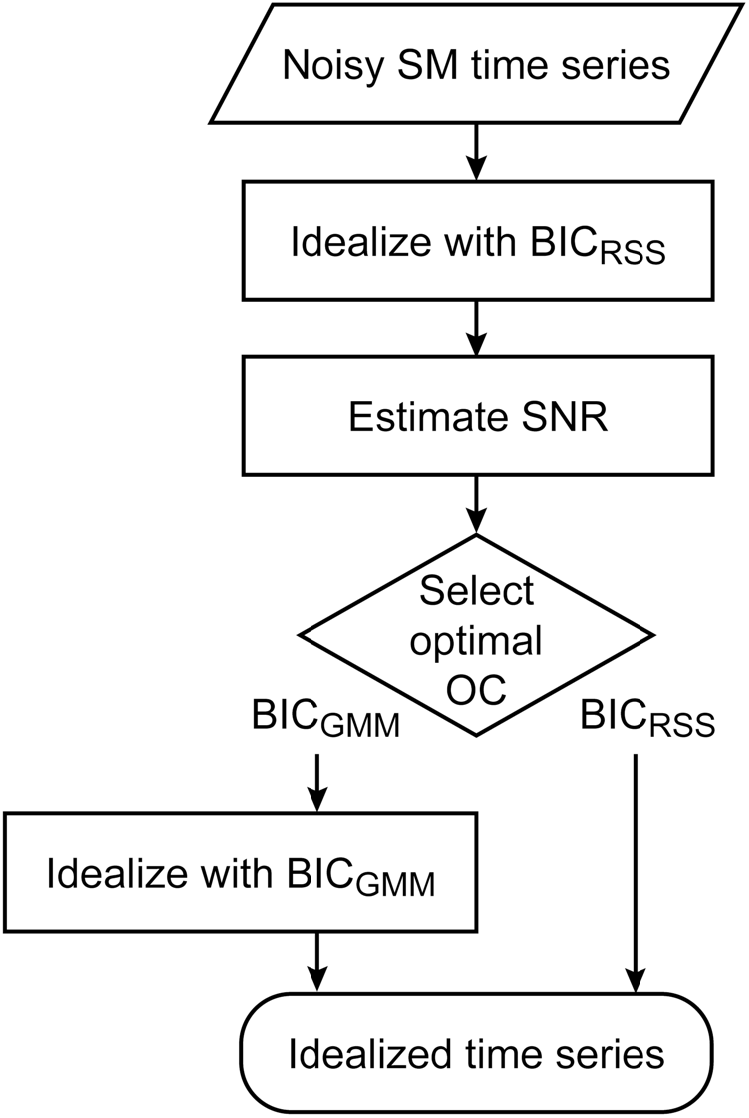
Workflow for unsupervised per-molecule optimal idealization by DISC.

## Discussion

The performance of the DISC algorithm with each of five different OCs (BIC_GMM_, BIC_RSS_, AIC, HQC, and MDL) was investigated for simulations reflecting typical SM fluorescence time series observations under variable experimental conditions. Each OC differentially impacts DISC’s performance for different conditions, with the SNR and number of samples in a time series being the primary determinants for the optimal OC. For nearly all conditions tested the OCs with similarly optimal F1 scores included either BIC_GMM_ or BIC_RSS_. BIC_RSS_ is typically optimal for time series with few samples and/or a low SNR, whereas BIC_GMM_ is typically optimal for time series with a thousand or more samples and high SNR. This difference in behavior can largely be attributed to the different penalty terms for each OC. Optimization of a variable penalty term does not generally outperform either BIC_GMM_ or BIC_RSS_, suggesting that a simple choice between the two OCs is sufficient for optimal performance by DISC in nearly all cases. Examination of the optimal OC under variable conditions (i.e. SNR and number of samples) reveal a general linear decision boundary that can be used to select the optimal OC for a given set of conditions. We further developed an estimator for the SNR of experimental SM time series that relates intensity level separation and Gaussian noise. Together, these developments establish a completely unsupervised workflow for the DISC algorithm that automates selection of the optimal OC on a per molecule basis.

Fluorescence imaging is a common approach for massively parallel observations of SM time series (Chen et al., 2014; Juette et al., 2016; Levene et al., 2003; Yan et al., 2018). Critically, intensity level separation versus noise ratios in SM fluorescence experimental datasets are often near our derived decision boundary, where optimal choice of OC is most impactful. Furthermore, stochastic fluorophore bleaching and nonuniformities in the optical pathways result in per-molecule variation in signal properties such as the observation time window and the ratio of the fluorescence intensity separation between distinct states to the noise within each state (Holden et al., 2010; Oostveldt et al., 1998). In addition to uncertainty regarding the underlying mechanism, per-molecule variation in state intensity levels challenges analysis with models such as HMMs having user-defined state intensity distributions. For such large datasets it is often the case that only a subset of molecules exhibit the behavior of interest. It is therefore ideal to avoid computationally intensive analyses such as HMMs on the potentially large fraction of non-relevant molecules. Rapid, unsupervised approaches are thus ideal in many cases, even if the result is only used to pre-screen a subset of molecules for further analysis, although these approaches can also be used for the full analysis.

DISC is a recently developed unsupervised approach for idealization of noisy SM time series that automates detection of the intensity levels within each individual time series without requiring postulation of a specific mechanistic model (White et al., 2020). Where possible, the additional constraint imposed by global optimization of a specific molecular mechanism or a singular set of state intensity distributions should be preferred. However, this approach is not appropriate for unknown mechanisms or experimental data with per-molecule variability in state intensity emissions. DISC not only handles these data efficiently but is also robust to within-state intensity fluctuations that can arise from changing orientation in polarized and/or exponentially decaying excitation fields or dye photodynamics (Dempsey et al., 2009; Levene et al., 2003; Stennett et al., 2015). Furthermore, DISC is orders of magnitude faster than other HMM or change-point methods while maintaining state-of-the-art accuracy, precision and recall (White et al., 2020). However, DISC relies on a user-specified OC to guide its idealization. Here, we show that the optimal choice of OC depends primarily on both the number of sample points and the signal and noise properties in each time series. Thus, maximizing the performance of DISC on experimental datasets with stochastic variability in these parameters requires selection of the optimal OC on a per-molecule basis.

By automating the per-molecule choice of OC we develop a fully unsupervised workflow that maximizes DISC’s performance across datasets with stochastic variability in observation conditions. As compared to DISC, the additional computational cost is either marginal or at most doubled owing to rerunning DISC with BIC_GMM_. Thus, this workflow does not negate one of DISC’s major advantages in that it is orders of magnitude faster than comparable approaches, and thus attractive for large high-throughput datasets. Given the prevalence of per-molecule stochastic variation in SM fluorescence observations and intensity level separation versus noise ratios that are often near our derived decision boundary, our approach provides an immediately useful tool to both optimize and speed their exploration and analysis, as well as that of other SM datasets exhibiting similar properties.

## Methods

### Simulations

All simulations and analyses were conducted in MATLAB version R2019a (Mathworks). SM time series were simulated as Markov chains of discrete dwells in distinct molecular states governed by the average rate of transitions between states (McKinney et al., 2006). The simulated Markov mechanisms are depicted in **Figure 1-figure supplement 1**. Ideal noiseless time series were constructed from the simulated state sequences by assigning an observable intensity to each state, after which Gaussian noise was added to generate noisy simulated observations. The standard deviation of the added noise *σ* was set to the ratio of the average separation between neighboring state intensity levels and the specified SNR of the simulation. For the 1-site model, state intensity levels were 0 and 1, so *σ* =1/*SNR*. Simulations for 2-site and 4-site models were generated by summing either two or four noisy 1-site simulations, respectively, consistent with observations of molecular association at multiple independent sites. For the three-state models, state intensity levels were 0.2, 0.6 and 0.8 to reflect typical smFRET observations, and *σ* = 0.3/*SNR*. All simulations have a uniform sample frequency *f*_*s*_, and transition rates between states are specified relative to the sample frequency. For the 1-site model, both forward and backward transitions rates were set to the same value which ranged from 0.01-1 *f*_*s*_. For the three-state models the fast rate *k*_*f*_ was set as described for the 1-site model and the slow rate *k*_*s*_ was set to 0.3*k*_*f*_. The fast rate always described transitions between the higher intensity levels. These prescriptions provide simulations that test performance on both equivalent and disparate rates within a given model. Where applicable we implemented state intensity heterogeneity on a per-event basis by adding a small stochastic offset to the mean intensity of individual events. The offset was drawn from an exponential distribution with a mean set to 4% of the state’s intensity level similar to prior observations of such fluctuations in measures of molecular association (White et al., 2020). For each model and each unique combination of SNR, number of samples per time series, and transition rate (*k* or *k*_*f*_), we simulated a total of 100,000 sample points split between 10-1000 time series depending on series length.

### Idealization performance

Noisy simulated time series were idealized with DISC using one of the flowing OCs: BIC_GMM_, BIC_RSS_, AIC, HQC, or MDL (Eqs. 2-10). In principle, a different OC could be selected for the segmentation and clustering portions of the DISC algorithm. Here, we only explore the impact of selecting a single OC for all portions of the algorithm. The general format for each OC is a summation of two terms that balance goodness of fit with a penalty for overfitting noise with overly complex models.

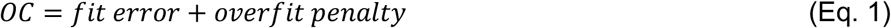

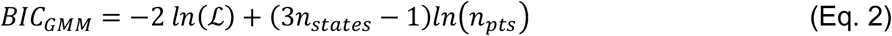

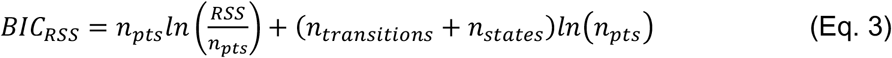

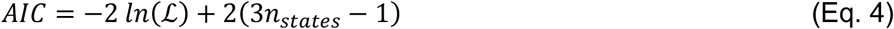

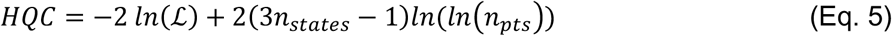

where ℒ is the likelihood for the estimated model defined as the product of likelihoods for each data point *y*(*t*_*i*_), each of which is described by a linear combination of likelihoods for each state’s Gaussian emission distribution 𝒩 with mean and standard deviation *μ*_*j*_ and *σ*_*j*_ and mixing coefficient *w*_*j*_ (Bishop, 2006).

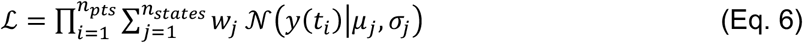

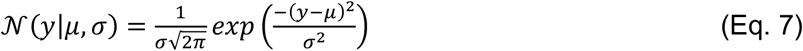

We define MDL as previously described by Shuang et al., 2014

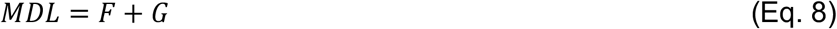

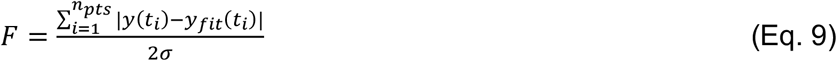

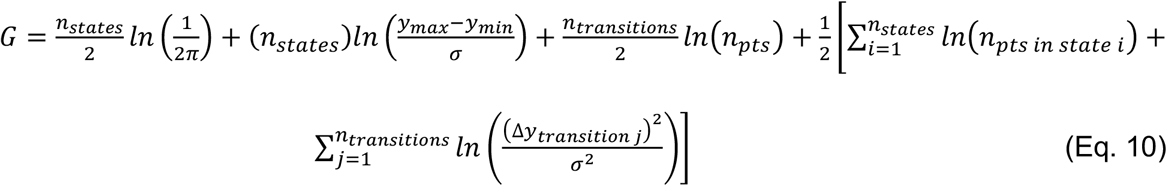

To quantify the quality of DISC’s idealizations, each event returned by DISC was determined to be either a true positive (TP), false positive (FP), or false negative (FN) based on the known simulated noiseless sequence. To prevent slight numerical differences between simulated and idealized intensity levels or subtle offsets in event onset or offset from being construed as errors, TP events were allowed to have intensities within ±10% of the known intensity level and onset or offset times within ±3 samples of the known event timing. Events classified as FPs were either extraneous events or correct events with the wrong intensity. FNs were defined as missed events. For each idealization, accuracy, precision, and recall were calculated as general performance metrics ranging from 0 (worst) to 1 (best) (Eqs. 11-13). The F1 score is a widely used overall metric for summarizing performance, and ranges from 0 (worst) to 1 (best) (Eq. 14) (Van Rijsbergen, 1979).

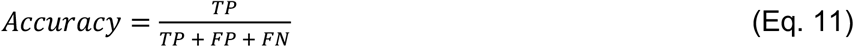

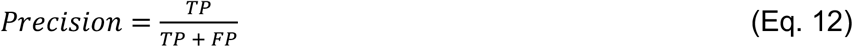

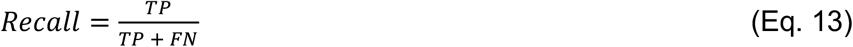

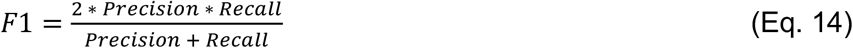

### OC penalty hyperparameter

To test the effects of a variable penalty term, we scaled the penalty terms of both BIC_GMM_ and BIC_RSS_ using a hyperparameter *λ* (see Eq. 15). For each unique combination of conditions (SNR, number of samples, transition rate), simulated traces from the 4-site model with added state intensity heterogeneity were idealized with DISC using either BIC_GMM_ or BIC_RSS_ for values of *λ* ranging from 0.001 to 10. The dependence of F1 scores on *λ* is shown in **Figure 1-figure supplements 9-10**. Notably, for nearly all conditions the optimal value of *λ* did not enhance performance beyond the better of either BIC_GMM_ or BIC_RSS_ with *λ* = 1.

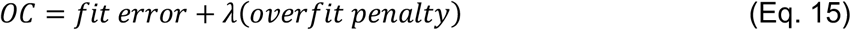

### Linear decision boundary for BIC_GMM_ vs BIC_RSS_

To generate a decision boundary for the optimal choice of either BIC_GMM_ or BIC_RSS_ based on the properties of the time series, we used MATLAB’s *fitcsvm* function from the *Statistics and Machine Learning Toolbox* (Mathworks) Every experimental condition (unique combination of SNR, number of samples and transition rate) for each tested model was labeled as either “BIC_GMM_ optimal” or “BIC_RSS_ optimal”. For each model, these labels were used to train a support vector machine (SVM) classifier to determine a linear decision boundary in the two-dimensional space of SNR and observation window (i.e. number of samples in a trace). This process was repeated for each model both with and without added state intensity heterogeneity **(Figure 2-figure supplement 1)**. To better visualize the extent to which BIC_GMM_ or BIC_RSS_ were optimal for a given experimental condition, we generated a preference score for each condition describing the fraction of simulations at that condition where BIC_GMM_ was optimal (Eq. 16). Additionally, decision-boundaries specific for each rate’s data were generated for all of the models with and without added state intensity heterogeneity. The lack of major shifts in boundary except at extreme transition rates (eg. transition rate approaching sampling rate) confirmed that a 2-dimensional boundary based on SNR and number of samples in the time series was appropriate **(Figure 2-figure supplement 2)**.

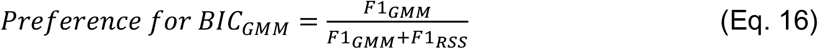

## Author Contributions

A.B. performed all simulations and analysis and contributed to writing the manuscript. M.P.G. conceived, designed and oversaw the project, and contributed to writing of the manuscript.

## Competing Interests

The authors declare no competing financial interests.

---

**Figure 1-figure supplement 1.**
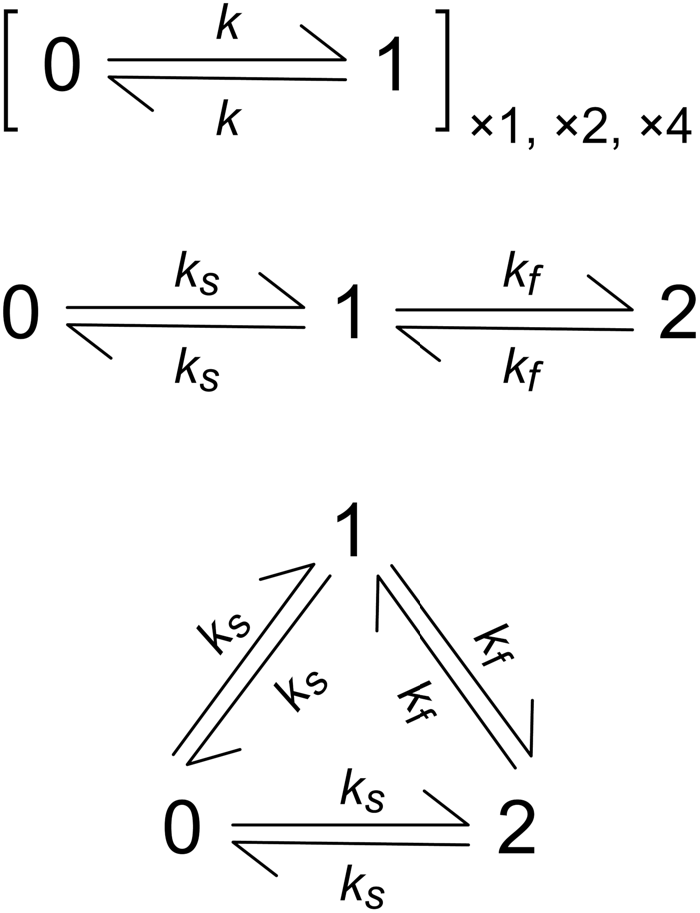
Markov models used for simulations. *top*) Two-state dynamics with equivalent forward and backward transition rates *k* for 1, 2 or 4 identical and independent sites. *middle and bottom*) Three-state dynamics with slow (*k*_*s*_) and fast (*k*_*f*_) transition rates.

**Figure 1-figure supplement 2.**
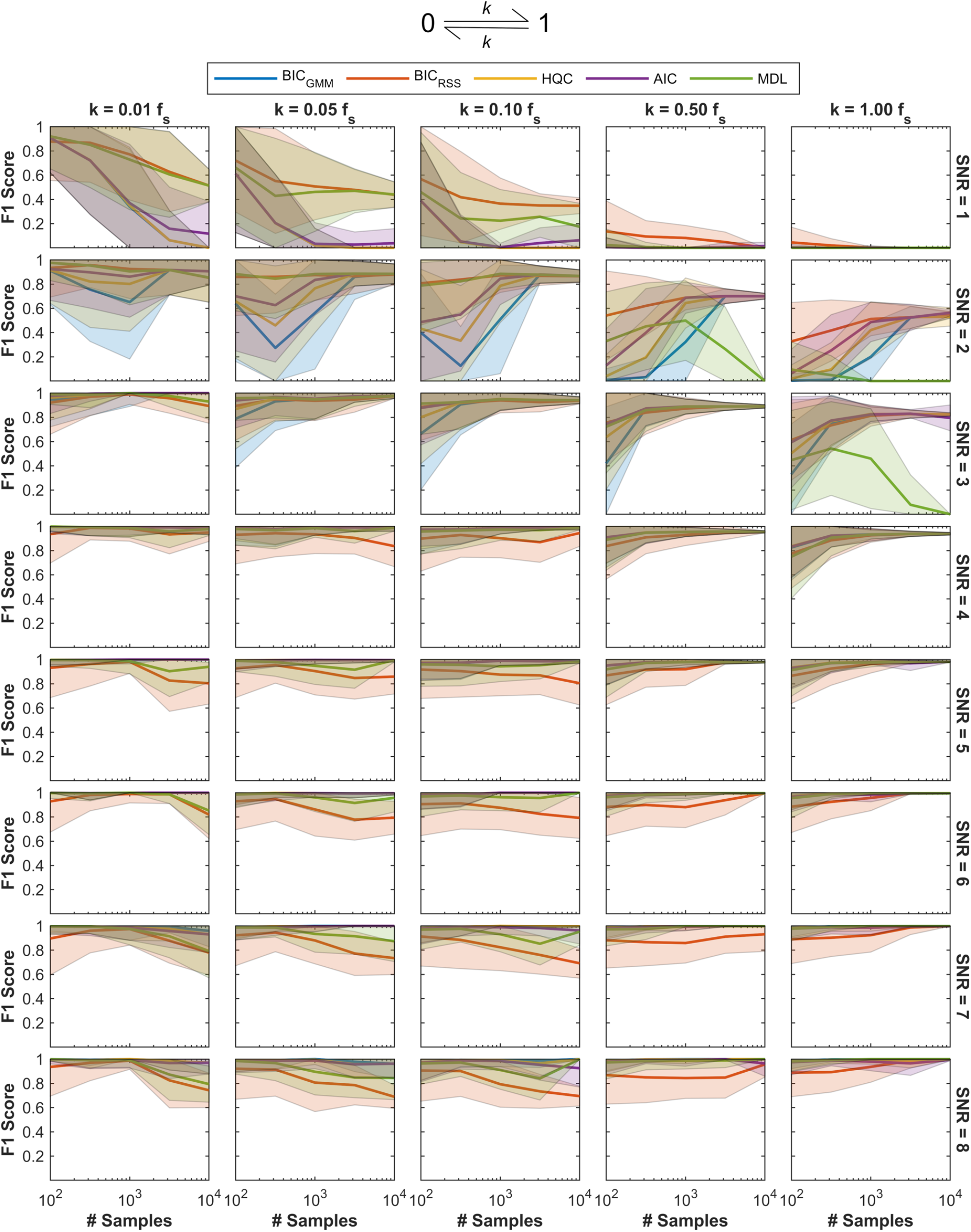
Impact of OC choice on DISC’s idealization performance. F1 score as a proxy for performance of DISC on idealization of noisy simulated SM time series for two-state dynamics (top) under various conditions (i.e. SNR, number of samples per trace, and transition rate *k* relative to the sample frequency *f*_*s*_) and with per-event state intensity heterogeneity. Performance with five different OCs are overlaid. Mean (line) and standard deviation (shaded region) for F1 scores (0-1: worst to perfect; Eq. 14) for 10-1000 simulated time series at each unique set of conditions (# samples and SNR) idealized with DISC using the indicated OC (see Methods).

**Figure 1-figure supplement 3.**
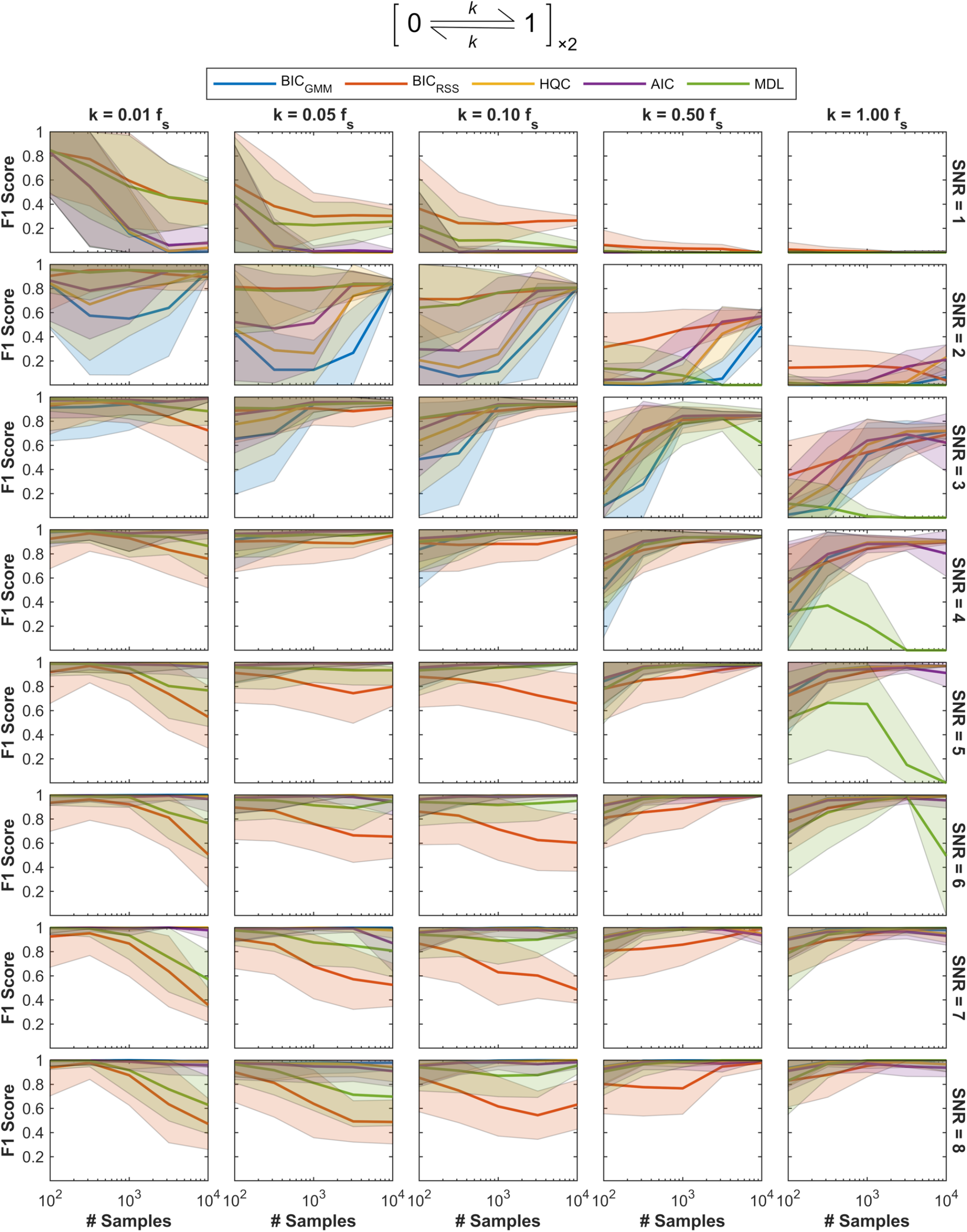
Impact of OC choice on DISC’s idealization performance. F1 score as a proxy for performance of DISC on idealization of noisy simulated SM time series for two-state dynamics at two identical and independent sites (top) under various conditions (i.e. SNR, number of samples per trace, and transition rate *k* relative to the sample frequency *f*_s_) and with per-event state intensity heterogeneity. Performance with five different OCs are overlaid. Mean (line) and standard deviation (shaded region) for F1 scores (0-1: worst to perfect; Eq. 14) for 10-1000 simulated time series at each unique set of conditions (# samples and SNR) idealized with DISC using the indicated OC (see Methods).

**Figure 1-figure supplement 4.**
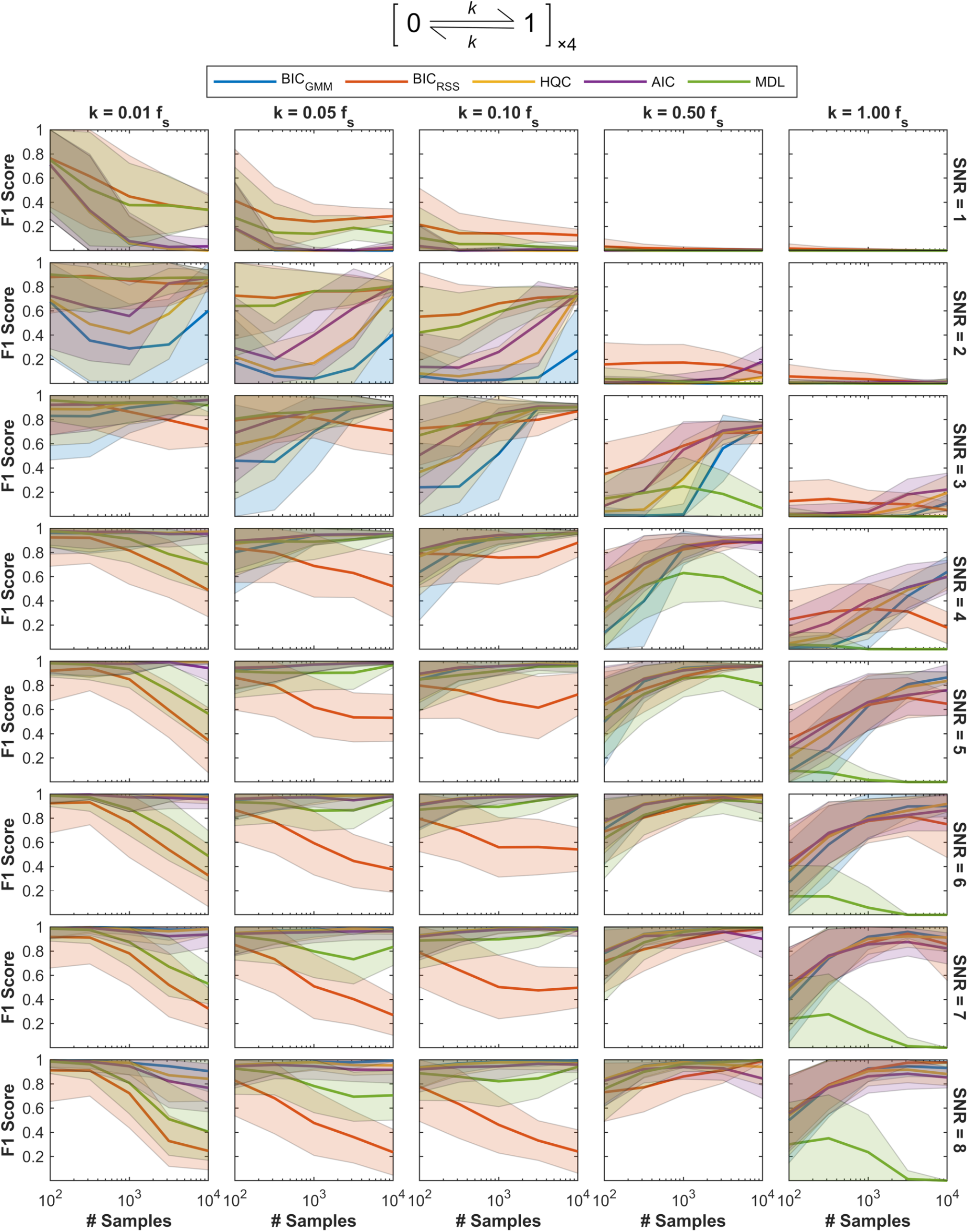
Impact of OC choice on DISC’s idealization performance. F1 score as a proxy for performance of DISC on idealization of noisy simulated SM time series for two-state dynamics at four identical and independent sites (top) under various conditions (i.e. SNR, number of samples per trace, and transition rate *k* relative to the sample frequency *f*_*s*_) and with per-event state intensity heterogeneity. Performance with five different OCs are overlaid. Mean (line) and standard deviation (shaded region) for F1 scores (0-1: worst to perfect; Eq. 14) for 10-1000 simulated time series at each unique set of conditions (# samples and SNR) idealized with DISC using the indicated OC (see Methods).

**Figure 1-figure supplement 5.**
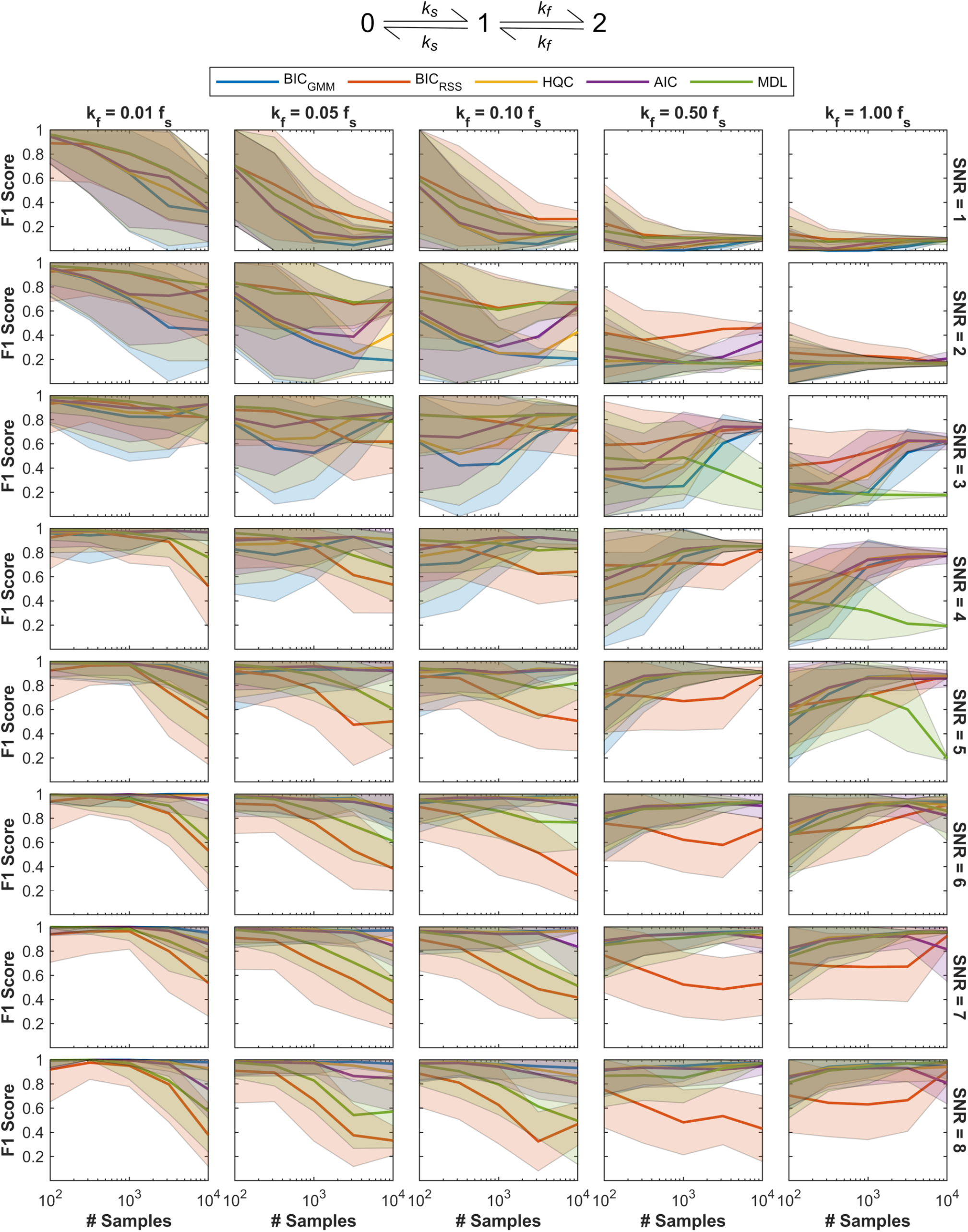
Impact of OC choice on DISC’s idealization performance. F1 score as a proxy for performance of DISC on idealization of noisy simulated SM time series for athree-state linear model (top) under various conditions (i.e. SNR, number of samples per trace, and transition rates *k*_*f*_ and *k*_*s*_ = 0.3*k*_*f*_ relative to the sample frequency *f*_*s*_) and with per-event state intensity heterogeneity. Performance with five different OCs are overlaid. Mean (line) and standard deviation (shaded region) for F1 scores (0-1: worst to perfect; Eq. 14) for 10-1000 simulated time series at each unique set of conditions (# samples and SNR) idealized with DISC using the indicated OC (see Methods).

**Figure 1-figure supplement 6.**
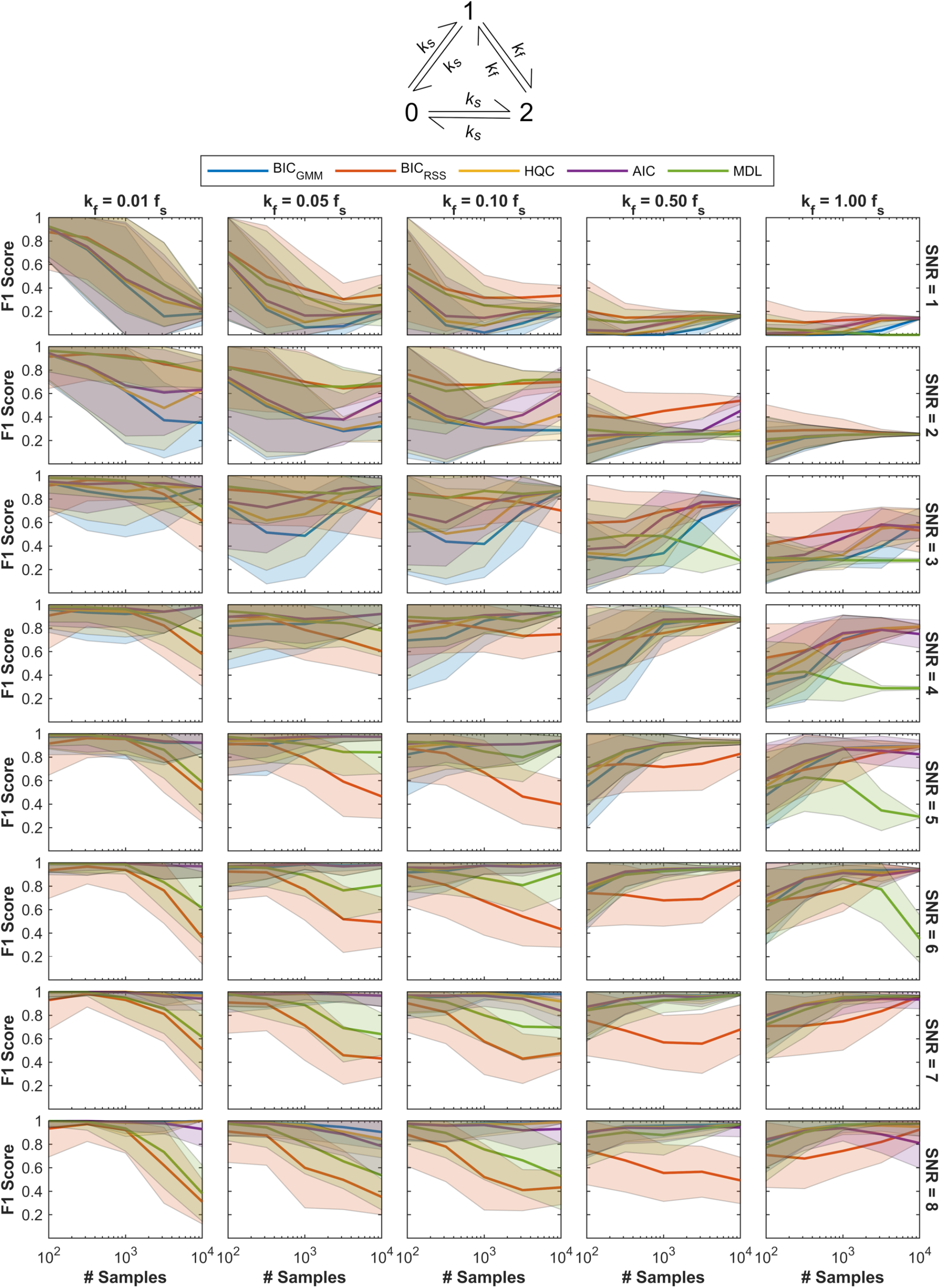
Impact of OC choice on DISC’s idealization performance. F1 score as a proxy for performance of DISC on idealization of noisy simulated SM time series for a three-state cyclic model (top) under various conditions (i.e. SNR, number of samples per trace, and transition rates *k*_*f*_ and *k*_*s*_ = 0.3*k*_*f*_ relative to the sample frequency *f*_*s*_) and with per-event state intensity heterogeneity. Performance with five different OCs are overlaid. Mean (line) and standard deviation (shaded region) for F1 scores (0-1: worst to perfect; Eq. 14) for 10-1000 simulated time series at each unique set of conditions (# samples and SNR) idealized with DISC using the indicated OC (see Methods).

**Figure 1-figure supplement 7.**
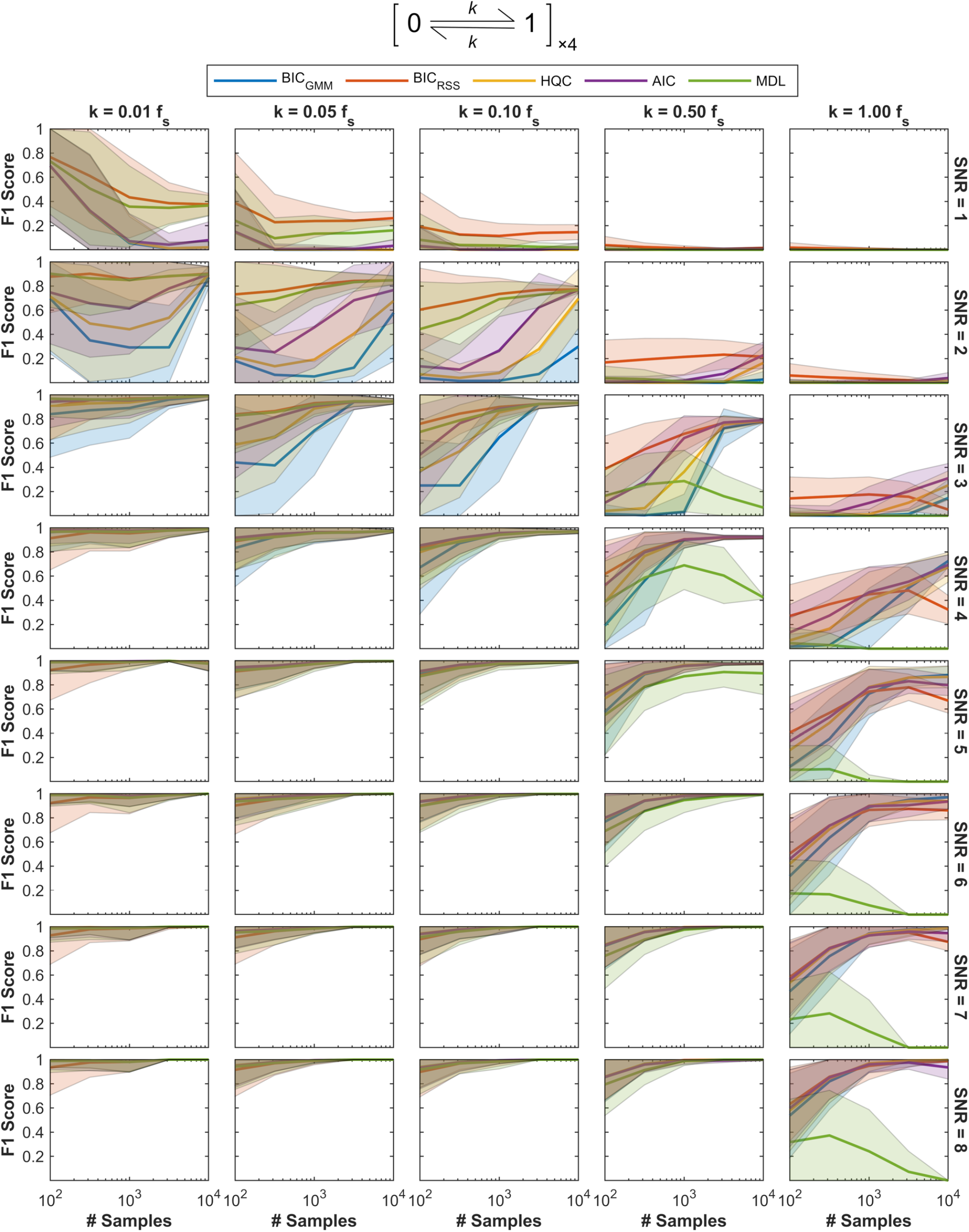
Impact of OC choice on DISC’s idealization performance. F1 score as a proxy for performance of DISC on idealization of noisy simulated SM time series for two-state dynamics at four identical and independent sites (top) under various conditions (i.e. SNR, number of samples per trace, and transition rate *k* relative to the sample frequency *f*_*s*_). No per-event state intensity heterogeneity was added. Performance with five different OCs are overlaid. Mean (line) and standard deviation (shaded region) for F1 scores (0-1: worst to perfect; Eq. 14) for 10-1000 simulated time series at each unique set of conditions (# samples and SNR) idealized with DISC using the indicated OC (see Methods).

**Figure 1-figure supplement 8.**
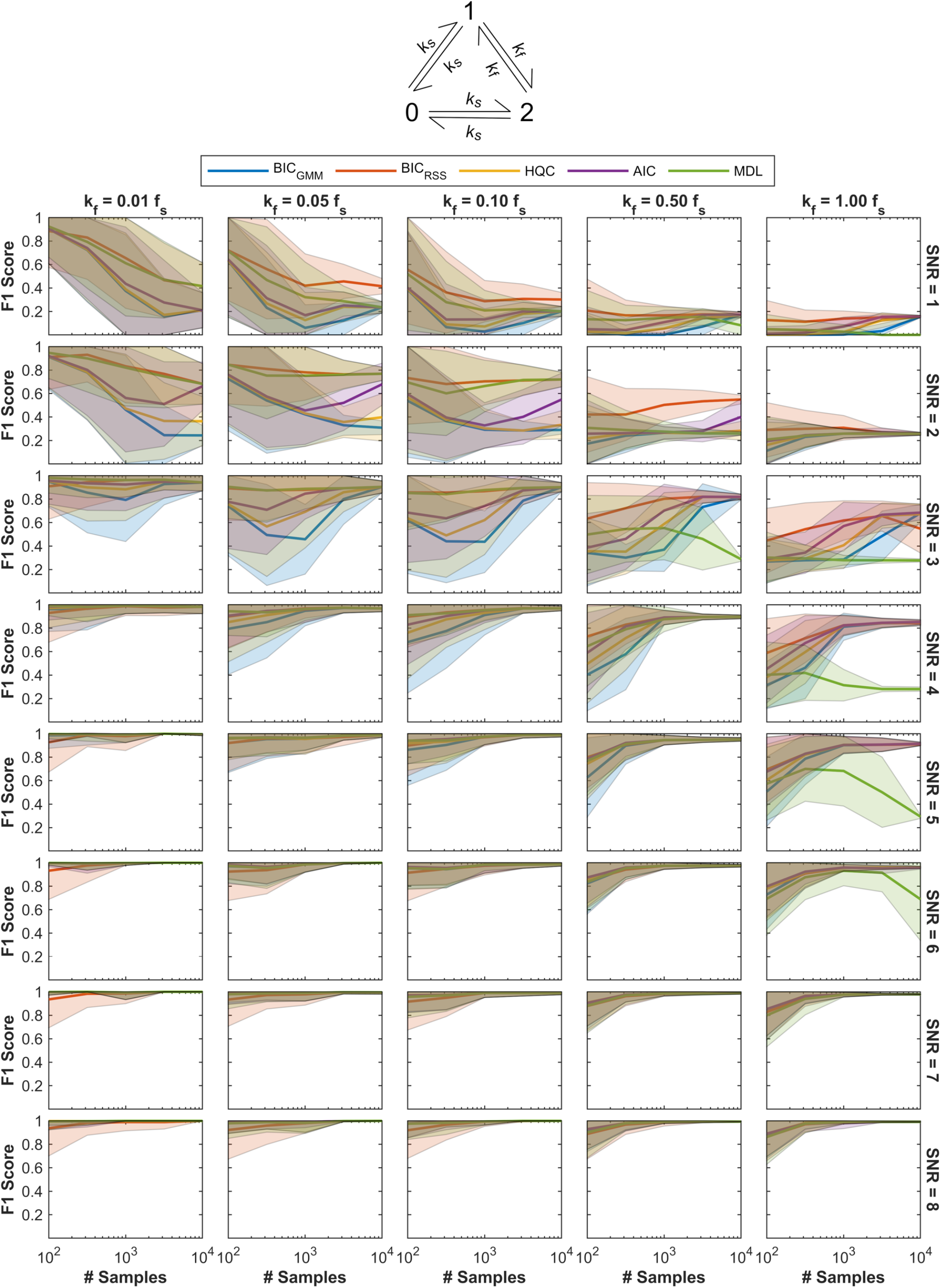
Impact of OC choice on DISC’s idealization performance. F1 score as a proxy for performance of DISC on idealization of noisy simulated SM time series for a three-state cyclic model (top) under various conditions (i.e. SNR, number of samples per trace, and transition rates *k*_*f*_ and *k*_*s*_ = 0.3*k*_*f*_ relative to the sample frequency *f*_*s*_). No per-event state intensity heterogeneity was added. Performance with five different OCs are overlaid. Mean (line) and standard deviation (shaded region) for F1 scores (0-1: worst to perfect; Eq. 14) for 10-1000 simulated time series at each unique set of conditions (# samples and SNR) idealized with DISC using the indicated OC (see Methods).

**Figure 1-figure supplement 9.**
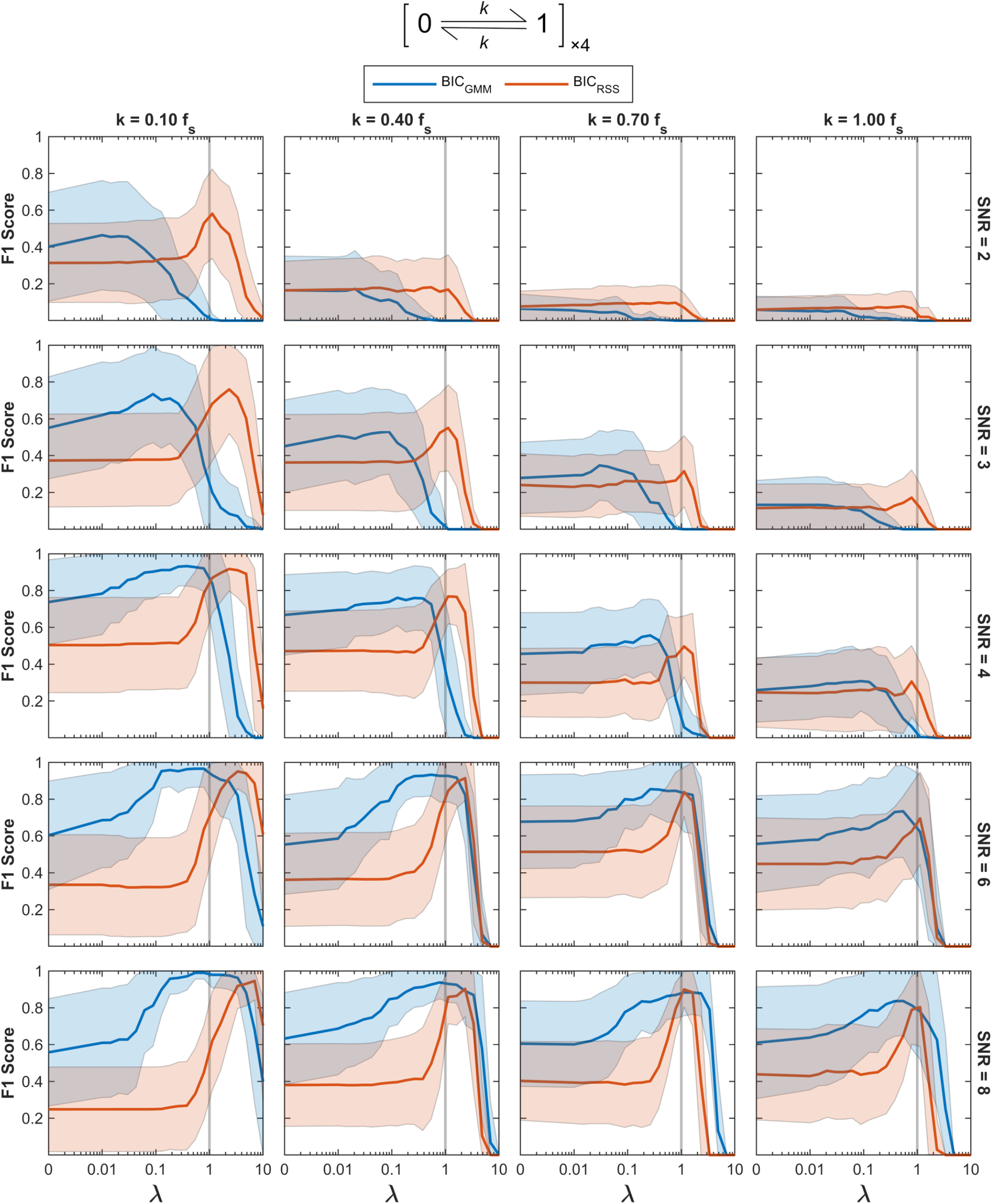
Impact of λ-scaled penalty terms on DISC’s idealization performance. F1 score as a proxy for performance of DISC on idealization of noisy simulated SM time series with 300 samples per trace for two-state dynamics at four identical and independent sites (top) under various conditions (i.e. SNR and transition rate *k* relative to the sample frequency*f*_*s*_) and with per-event state intensity heterogeneity. Performance with BIC_GMM_ and BIC_RSS_ overlaid with vertical line indicating λ = 1. Mean (line) and standard deviation (shaded region) for F1 scores (0-1: worst to perfect; Eq. 14) for 10-1000 simulated time series at each unique set of conditions (# samples and SNR).

**Figure 1-figure supplement 10.**
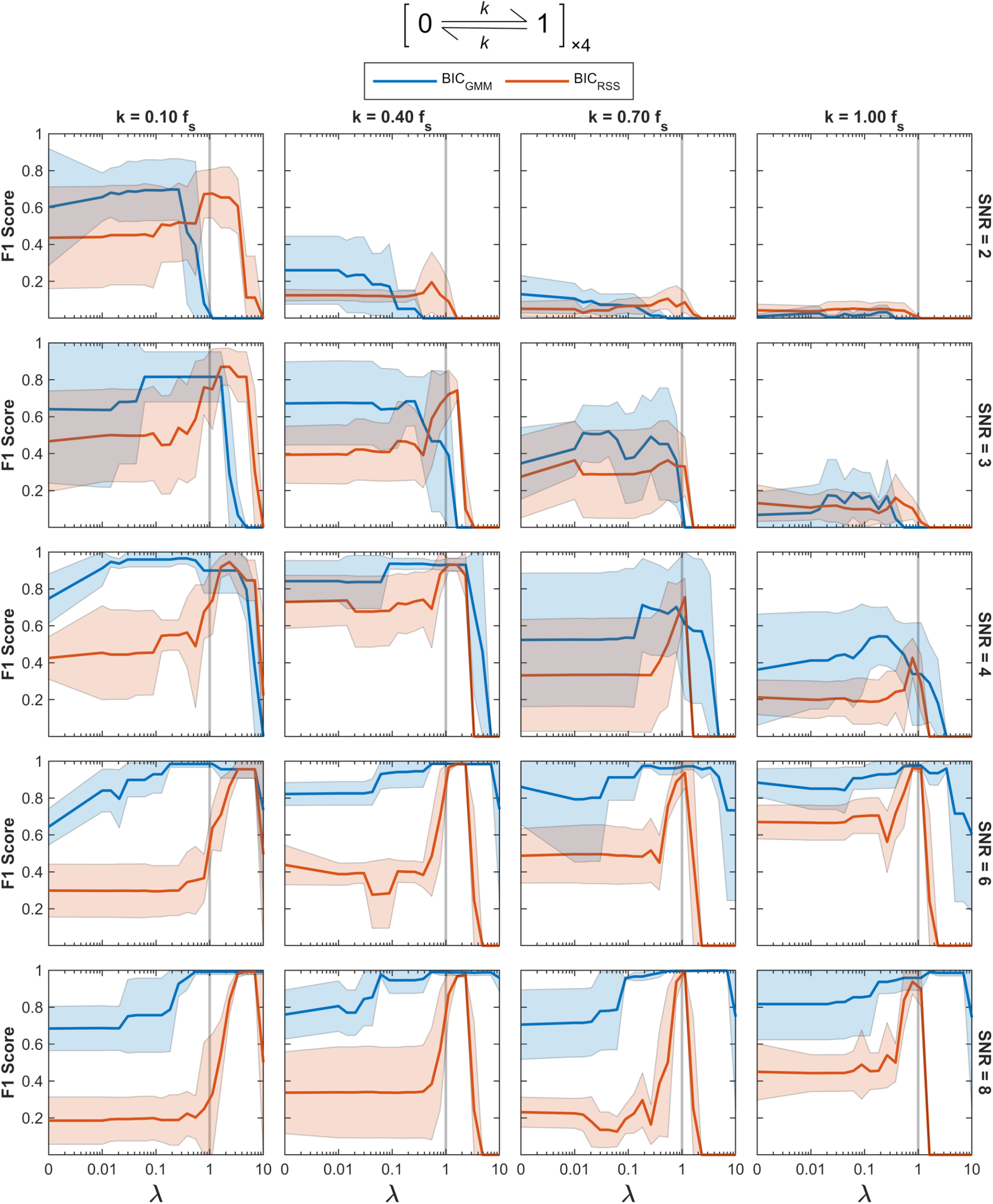
Impact of λ-scaled penalty terms on DISC’s idealization performance. F1 score as a proxy for performance of DISC on idealization of noisy simulated SM time series with 3000 samples per trace for two-state dynamics at four identical and independent sites (top) under various conditions (i.e. SNR and transition rate *k* relative to the sample frequency *f*_*s*_) and with per-event state intensity heterogeneity. Performance with BIC_GMM_ and BIC_RSS_ overlaid with vertical line indicating λ = 1. Mean (line) and standard deviation (shaded region) for F1 scores (0-1: worst to perfect; Eq. 14) for 10-1000 simulated time series at each unique set of conditions (# samples and SNR).

**Figure 2-figure supplement 1.**
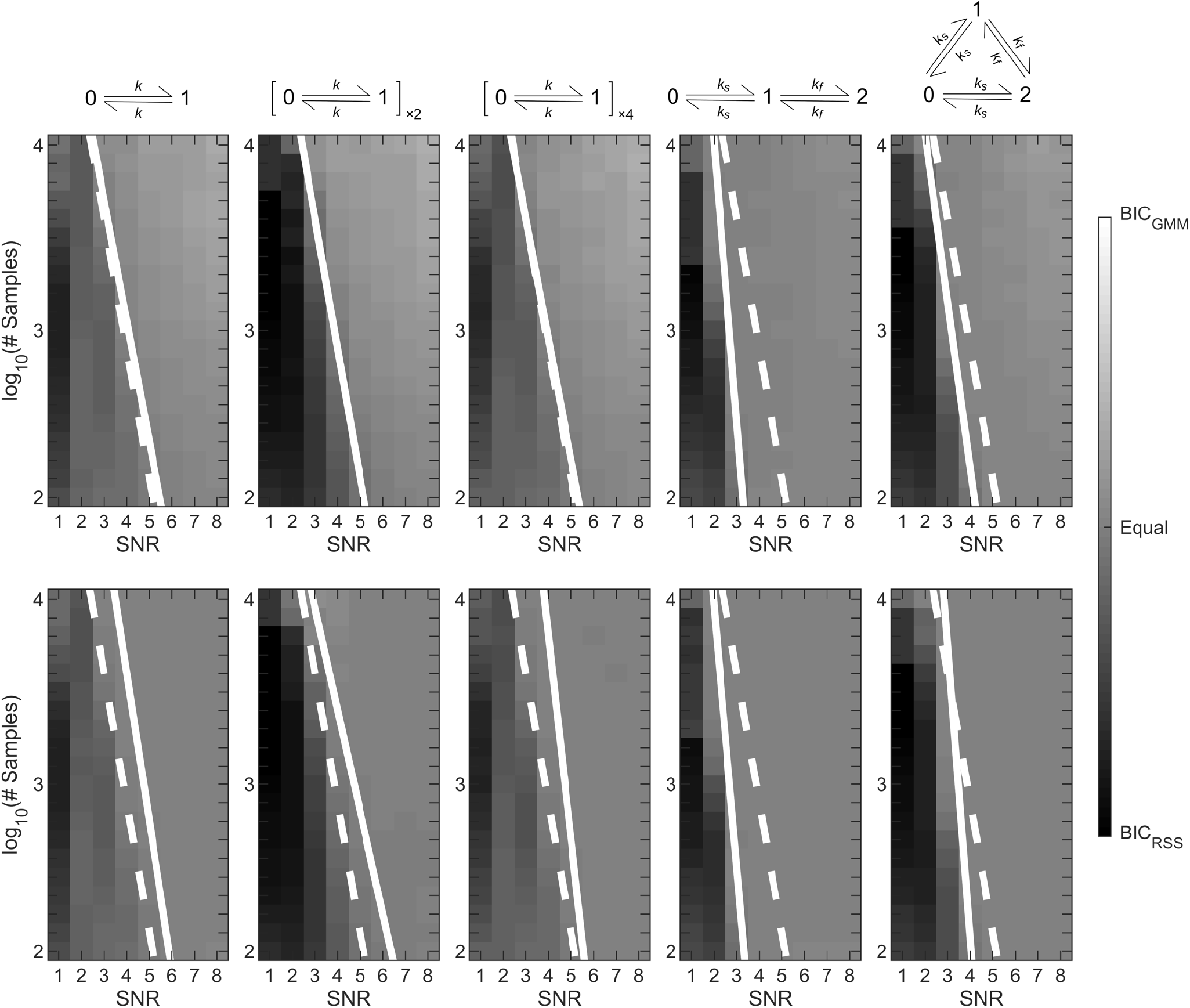
Linear decision boundaries. Heatmaps indicating preference for BIC_RSS_ (black) or BIC_GMM_ (white) for 1000 simulations at each unique combination of conditions (SNR and number of samples per trace). Each column represents simulations for one of the five Markov models depicted at the top of the column (from left to right: two-state dynamics, two-state dynamics at two independent sites, two-state dynamics at four independent sites, three-state linear, and three-state cyclic) with (top row) and without (bottom row) per-event state intensity heterogeneity. Solid white lines are the linear decision boundary determined by an SVM classifier. The dashed white line is the decision boundary for the model with two-state dynamics at four identical and independent sites including state intensity heterogeneity.

**Figure 2-figure supplement 2.**
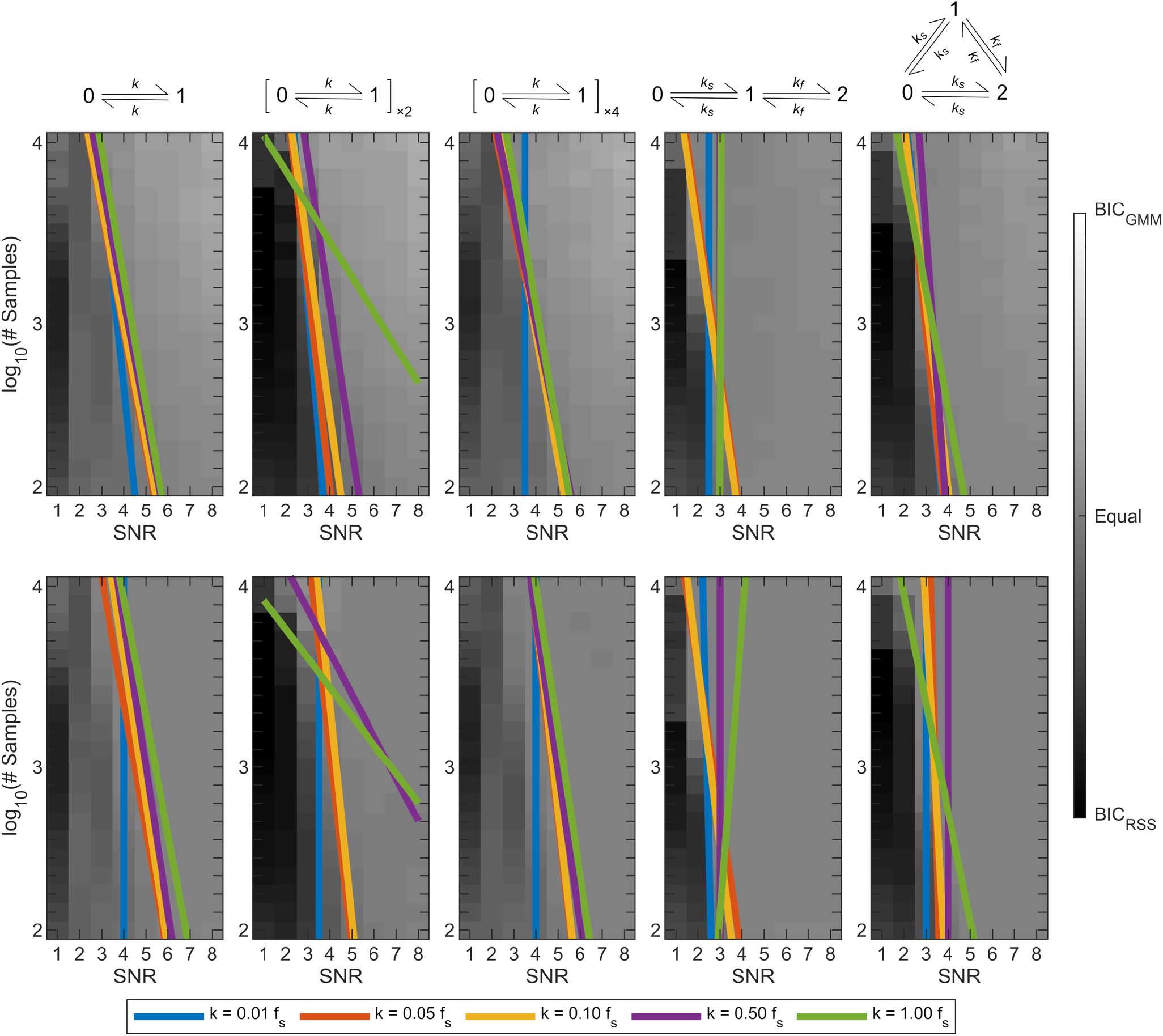
Dependence of linear decision boundaries on transition rates. Heatmaps indicating preference for BIC_RSS_ (black) or BIC_GMM_ (white) for 1000 simulations at each unique combination of conditions (SNR, number of samples per trace, transition rate) averaged across transition rates. Each column represents simulations for one of the five Markov models depicted at the top of the column (from left to right: two-state dynamics, two-state dynamics at two independent sites, two-state dynamics at four independent sites, three-state linear, and three-state cyclic) with (top row) and without (bottom row) per-event state intensity heterogeneity. Colored lines are the linear decision boundary determined by an SVM classifier for various transition rates relative to the sample frequency *f*_*s*_.

## Notes

https://github.com/marcel-goldschen-ohm/DISC

### Competing Interest Statement

The authors have declared no competing interest.

## References

Akaike, H. (1974). A new look at the statistical model identification. IEEE Transactions on Automatic Control, 19(6), 716–723. https://doi.org/10.1109/TAC.1974.1100705

Altman, R. B., Terry, D. S., Zhou, Z., Zheng, Q., Geggier, P., Kolster, R. A., Zhao, Y., Javitch, J. A., Warren, J. D., & Blanchard, S. C. (2012). Cyanine fluorophore derivatives with enhanced photostability. Nature Methods, 9(1), 68–71. https://doi.org/10.1038/nmeth.1774

Bishop, C. (2006). Pattern Recognition and Machine Learning. Springer.

Blanco, M., Johnson-Buck, A., & Walter, N. (2013). Hidden Markov Modeling in Single-Molecule Biophysics. /paper/Hidden-Markov-Modeling-in-Single-Molecule-Blanco-Johnson-Buck/31213d29942de2750cbd6a214b4086eb015a1acf

Blanco, Mario, & Walter, N. G. (2010). Analysis of complex single-molecule FRET time trajectories. Methods in Enzymology, 472, 153–178. https://doi.org/10.1016/S0076-6879(10)72011-5

Bronson, J. E., Fei, J., Hofman, J. M., Gonzalez, J., & Wiggins, C. H. (2009). Learning Rates and States from Biophysical Time Series: A Bayesian Approach to Model Selection and Single-Molecule FRET Data. Biophysical Journal, 97(12), 3196–3205. https://doi.org/10.1016/j.bpj.2009.09.031

Celik, N., O’Brien, F., Brennan, S., Rainbow, R. D., Dart, C., Zheng, Y., Coenen, F., & Barrett-Jolley, R. (2020). Deep-Channel uses deep neural networks to detect single-molecule events from patch-clamp data. Communications Biology, 3(1), 1–10. https://doi.org/10.1038/s42003-019-0729-3

Chen, J., Dalal, R. V., Petrov, A. N., Tsai, A., O’Leary, S. E., Chapin, K., Cheng, J., Ewan, M., Hsiung, P.-L., Lundquist, P., Turner, S. W., Hsu, D. R., & Puglisi, J. D. (2014). High-throughput platform for real-time monitoring of biological processes by multicolor single-molecule fluorescence. Proceedings of the National Academy of Sciences, 111(2), 664– 669. https://doi.org/10.1073/pnas.1315735111

Dempsey, G. T., Bates, M., Kowtoniuk, W. E., Liu, D. R., Tsien, R. Y., & Zhuang, X. (2009). Photoswitching Mechanism of Cyanine Dyes. Journal of the American Chemical Society, 131(51), 18192–18193. https://doi.org/10.1021/ja904588g

Goldschen-Ohm, M. P., Klenchin, V. A., White, D. S., Cowgill, J. B., Cui, Q., Goldsmith, R. H., & Chanda, B. (2016). Structure and dynamics underlying elementary ligand binding events in human pacemaking channels. ELife, 5, e20797. https://doi.org/10.7554/eLife.20797

Greenfeld, M., Pavlichin, D. S., Mabuchi, H., & Herschlag, D. (2012). Single Molecule Analysis Research Tool (SMART): An Integrated Approach for Analyzing Single Molecule Data. PLoS ONE, 7(2). https://doi.org/10.1371/journal.pone.0030024

Halabi, E. A., Pinotsi, D., & Rivera-Fuentes, P. (2019). Photoregulated fluxional fluorophores for live-cell super-resolution microscopy with no apparent photobleaching. Nature Communications, 10(1), 1232. https://doi.org/10.1038/s41467-019-09217-7

Hannan, E. J., & Quinn, B. G. (1979). The Determination of the Order of an Autoregression. Journal of the Royal Statistical Society. Series B (Methodological), 41(2), 190–195.

Hines, K. E., Bankston, J. R., & Aldrich, R. W. (2015). Analyzing Single-Molecule Time Series via Nonparametric Bayesian Inference. Biophysical Journal, 108(3), 540–556. https://doi.org/10.1016/j.bpj.2014.12.016

Holden, S. J., Uphoff, S., Hohlbein, J., Yadin, D., Le Reste, L., Britton, O. J., & Kapanidis, A. N. (2010). Defining the Limits of Single-Molecule FRET Resolution in TIRF Microscopy. Biophysical Journal, 99(9), 3102–3111. https://doi.org/10.1016/j.bpj.2010.09.005

Juette, M. F., Terry, D. S., Wasserman, M. R., Altman, R. B., Zhou, Z., Zhao, H., & Blanchard, S. C. (2016). Single-molecule imaging of non-equilibrium molecular ensembles on the millisecond timescale. Nature Methods, 13(4), 341–344. https://doi.org/10.1038/nmeth.3769

Kadane, J. B., & Lazar, N. A. (2004). Methods and Criteria for Model Selection. Journal of the American Statistical Association, 99(465), 279–290.

Levene, M. J., Korlach, J., Turner, S. W., Foquet, M., Craighead, H. G., & Webb, W. W. (2003). Zero-Mode Waveguides for Single-Molecule Analysis at High Concentrations. Science, 299(5607), 682–686. https://doi.org/10.1126/science.1079700

Li, J., Zhang, L., Johnson-Buck, A., & Walter, N. G. (2020). Automatic classification and segmentation of single-molecule fluorescence time traces with deep learning. Nature Communications, 11(1), 5833. https://doi.org/10.1038/s41467-020-19673-1

McKinney, S. A., Joo, C., & Ha, T. (2006). Analysis of Single-Molecule FRET Trajectories Using Hidden Markov Modeling. Biophysical Journal, 91(5), 1941–1951. https://doi.org/10.1529/biophysj.106.082487

Miller, H., Zhou, Z., Shepherd, J., Wollman, A. J. M., & Leake, M. C. (2018). Single-molecule techniques in biophysics: A review of the progress in methods and applications. Reports on Progress in Physics. Physical Society (Great Britain), 81(2), 024601. https://doi.org/10.1088/1361-6633/aa8a02

Oostveldt, P. V., Verhaegen, F., & Messens, K. (1998). Heterogeneous photobleaching in confocal microscopy caused by differences in refractive index and excitation mode. Cytometry, 32(2), 137–146. https://doi.org/10.1002/(SICI)1097-0320(19980601)32:2<137::AID-CYTO9>3.0.CO;2-I

Priestley, M. B. (2004). Spectral analysis and time series (Repr). Elsevier.

Schwarz, G. (1978). Estimating the Dimension of a Model. Annals of Statistics, 6(2), 461–464. https://doi.org/10.1214/aos/1176344136

Sgouralis, I., & Pressé, S. (2017). An Introduction to Infinite HMMs for Single-Molecule Data Analysis. Biophysical Journal, 112(10), 2021–2029. https://doi.org/10.1016/j.bpj.2017.04.027

Shuang, B., Cooper, D., Taylor, J. N., Kisley, L., Chen, J., Wang, W., Li, C. B., Komatsuzaki, T., & Landes, C. F. (2014). Fast Step Transition and State Identification (STaSI) for Discrete Single-Molecule Data Analysis. The Journal of Physical Chemistry Letters, 5(18), 3157– 3161. https://doi.org/10.1021/jz501435p

Soltani, M., Lin, J., Forties, R. A., Inman, J. T., Saraf, S. N., Fulbright, R. M., Lipson, M., & Wang, M. D. (2014). Nanophotonic trapping for precise manipulation of biomolecular arrays. Nature Nanotechnology, 9(6), 448–452. https://doi.org/10.1038/nnano.2014.79

Stennett, E. M. S., Ciuba, M. A., Lin, S., & Levitus, M. (2015). Demystifying PIFE: The Photophysics Behind the Protein-Induced Fluorescence Enhancement Phenomenon in Cy3. The Journal of Physical Chemistry Letters, 6(10), 1819–1823. https://doi.org/10.1021/acs.jpclett.5b00613

Van Rijsbergen, C. J. (1979). Information Retrieval (2nd ed.). Butterworths. http://www.dcs.gla.ac.uk/Keith/Chapter.7/Ch.7.html#fn0

White, D. S., Goldschen-Ohm, M. P., Goldsmith, R. H., & Chanda, B. (2020). Top-down machine learning approach for high-throughput single-molecule analysis. ELife, 9, e53357. https://doi.org/10.7554/eLife.53357

Xiaobo Zhou, & Wong, S.T. C. (2006). Informatics challenges of high-throughput microscopy. IEEE Signal Processing Magazine, 23(3), 63–72. https://doi.org/10.1109/MSP.2006.1628879

Xu, J., Qin, G., Luo, F., Wang, L., Zhao, R., Li, N., Yuan, J., & Fang, X. (2019). Automated Stoichiometry Analysis of Single-Molecule Fluorescence Imaging Traces via Deep Learning. Journal of the American Chemical Society, 141(17), 6976–6985. https://doi.org/10.1021/jacs.9b00688

Yan, R., Moon, S., Kenny, S. J., & Xu, K. (2018). Spectrally Resolved and Functional Super-resolution Microscopy via Ultrahigh-Throughput Single-Molecule Spectroscopy. Accounts of Chemical Research, 51(3), 697–705. https://doi.org/10.1021/acs.accounts.7b00545

